# Sensitisation of colonic nociceptors by IL-13 is dependent on JAK and p38 MAPK activity

**DOI:** 10.1101/2022.10.26.511877

**Authors:** Katie H. Barker, James P. Higham, Luke A. Pattison, Iain P. Chessell, Fraser Welsh, Ewan St. J. Smith, David C. Bulmer

## Abstract

The effective management of visceral pain is a significant unmet clinical need for those affected by gastrointestinal diseases, such as inflammatory bowel disease (IBD). The rational design of novel analgesics requires a greater understanding of the mediators and mechanisms underpinning visceral pain. Interleukin-13 (IL-13) production by immune cells residing in the gut is elevated in IBD, and IL-13 appears to be important in the development of experimental colitis. What’s more, receptors for IL-13 are expressed by neurons innervating the colon, though it is not known whether IL-13 plays any role in visceral nociception *per se*. To resolve this, we employed Ca^2+^ imaging of cultured sensory neurons and *ex vivo* electrophysiological recording from the lumbar splanchnic nerve innervating the distal colon. Ca^2+^ imaging revealed the stimulation of small-diameter, capsaicin-sensitive sensory neurons by IL-13, indicating that IL-13 likely stimulates nociceptors. IL-13-evoked Ca^2+^ signals were attenuated by inhibition of Janus (JAK) and p38 kinases. In the lumbar splanchnic nerve, IL-13 did not elevate baseline firing, nor sensitise the response to capsaicin application, but did enhance the response to distention of the colon. In line with Ca^2+^ imaging experiments, IL-13-mediated sensitisation of the afferent response to colon distention was blocked by inhibition of either JAK or p38 kinase signalling. Together, these data highlight a potential role for IL-13 in visceral nociception and implicate JAK and p38 kinases in pro-nociceptive signalling downstream of IL-13.

**Key Points Summary:** The pro-inflammatory cytokine IL-13 is elevated in gastrointestinal (GI) diseases and known to sensitise sensory neurons. This study confirms a role for IL-13 in colonic afferent sensitisation and defines a role for downstream JAK and p38 MAPK signalling in colonic mechanosensitisation.

IL-13-mediated increase in [Ca^2+^]_i_ in capsaicin-sensitive sensory neurons is dependent on p38 MAPK and JAK signalling.

IL-13-induced sensitisation of colonic afferents to noxious mechanical distension is abolished by inhibition of p38 MAPK and JAK.

We have built on the current understanding of IL-13 and its neuronal interactions, highlighting the therapeutic potential of targeting p38 MAPK and JAK signalling pathways to treat visceral pain in GI disease.

## Introduction

Pro-inflammatory cytokines can directly stimulate and/or sensitise dorsal root ganglion (DRG) neurons, as well as sensitising colonic afferent responses to noxious stimuli, thereby implicating these mediators in hypersensitivity and pain in GI disease. Whilst much research has already been conducted to determine the role of cytokines like tumour necrosis factor α (TNFα) in pain signalling pathways, some mediators remain less studied. One such cytokine is interleukin-13 (IL-13), a member of the T helper (Th) 2 family of cytokines, alongside IL-3, IL-4, IL-5, and IL-9. In humans, IL-13 is a 10-14 kDa immunoregulatory protein produced by activated CD4^+^ T cells and immune cells like mast cells, eosinophils, and basophils (Smithgall et al., 2008; Toru et al., 1998; Woerly et al., 2002). The gene encoding IL-13 in humans is located on chromosome 5q31 and consists of four exons and three introns, just upstream of the gene encoding IL-4 (Frazer et al., 1997; Iwaszko et al., 2021). Although IL-13 and IL-4 only share 25% similarity at the amino acid level, the two cytokines share overlapping roles in allergic inflammation and the response to parasite infection (Finkelman et al., 2004; Gour & Wills-Karp, 2015; Wills-Karp et al., 1998).

Given the overlapping physiological roles of IL-4 and IL-13, it should perhaps be no surprise that they also share similar signalling pathways, with both cytokines activating the heterodimeric type II receptor, which consists of a 140 kDa IL-4Rα and a 65-70 kDa IL-13Rα1 subunit (Aman et al., 1996; Khurana Hershey, 2003). IL-13Rα1 monomers bind IL-13 with low affinity, but dimerization with IL-4Rα increases binding affinity and receptor functionality (McKenzie et al., 1993; Miloux et al., 1997). Whilst the IL-4Rα subunit is also found together with the γc subunit in the type I receptor that interacts exclusively with IL-4 (Zurawski et al., 1995), IL-13 similarly binds with higher affinity to the IL-13Rα2 receptor, which is often referred to as a decoy receptor owing to a lack of signal transduction following IL-13 binding (Khurana Hershey, 2003).

The different receptor subunits of IL-4 and IL-13 receptors (other than IL-13Rα) activate common downstream signalling pathways upon cytokine binding. For example, the IL-4Rα receptor complexes are closely associated with Janus kinase (JAK) proteins; JAK1 associates with the IL-4Rα chain, JAK2/tyrosine kinase 2 (Tyk2) with IL-13Rα1, and JAK3 with γc (Goenka & Kaplan, 2011; Junttila, 2018; Kelly-Welch et al., 2003). JAK proteins phosphorylate IL-4Rα and recruit signal transducer and activator of transcription 6 (STAT6) (Nelms et al., 1999; Takeda et al., 1996). Once phosphorylated, STAT6 translocates to the nucleus (Goenka & Kaplan, 2011; Junttila, 2018; Kelly-Welch et al., 2003), but this interaction may also lead to activation of phosphatidylinositol 3-kinase (PI3K) and MAPK signalling pathways, as demonstrated in a range of *in vitro* systems (Atherton et al., 2003; Hecker et al., 2010; Iwashita et al., 2003; Wright et al., 1999).

IL-13 production is elevated in gastrointestinal (GI) diseases, with increased production by natural killer T (NKT) cells (Fuss et al., 2004, 2014). Moreover, other cell types within the gut, for example innate lymphoid cells (ILCs) and mast cells, can be stimulated to produce IL-13 by gut epithelium-derived cytokines, such as IL-33 and IL-25, which themselves are produced during inflammation and infection (Beltrán et al., 2010; Pastorelli et al., 2010; Tanaka et al., 2010; Zhao et al., 2010). Increased secretion of IL-13 via this mechanism may in part induce and maintain chronic idiopathic inflammation within the GI tract.

In experimental mouse models of colitis, IL-4 and IL-13 production contributes to disease pathogenesis with levels increasing progressively from pre- to early- to late-stage disease (Spencer et al., 2002). Moreover, neutralisation of IL-13 by IL-13Rα2-Fc administration (Heller et al., 2002) or use of a bifunctional therapeutic targeting IL-4 and IL-13 (Kasaian et al., 2014) prevented oxazolone-induced colitis in mouse models. In dextran sodium sulphate (DSS)-induced colitis, IL-13 secretion is also upregulated (Shajib et al., 2013) and disease severity has also been linked to mucosal levels of IL-13 (Okamura et al., 2014).

In inflammatory bowel disease (IBD) patients however, the role of IL-13 is less clear, with contrasting results reported from patient samples. Early studies reported reduced IL-13 concentrations in the inflamed mucosa of ulcerative colitis (UC) patients compared to active Crohn’s disease (CD) and healthy controls (Vainer et al., 2000) and more recently this has been reinforced by data showing similar levels of mucosal IL-13 mRNA and IL-13 protein production in active UC, inactive UC and healthy controls (Biancheri et al., 2014). However, *ex vivo* stimulation of lamina propria T cells taken from resected specimens of UC patients produced increased protein levels of IL-13 compared to stimulation of cells isolated from CD and healthy individuals (Fuss et al., 2004; Heller et al., 2005). In support of this, increased levels of IL-13Rα1 are present in *ex vivo* intestinal epithelial cells isolated from UC patients (Mandal & Levine, 2010) and increased expression of phosphorylated STAT6 has been identified in active UC with respect to CD and healthy controls (Rosen et al., 2011). Although other signalling pathways may activate STAT6 (Patel et al., 1996), IL-4 and IL-13 are by far the primary drivers of STAT6 phosphorylation (Goenka & Kaplan, 2011).

Thus far, IL-13 has been identified as a key driver in allergic asthma and has therefore influenced the production of therapeutics targeting the IL-13 pathway, including antibodies like anrukinzumab, lebrikizumab and tralokinumab, which prevent IL-13 signalling through IL-13Rα1 and IL-13Rα2, whilst maintaining IL-4 signalling through the type I receptor (May et al., 2012). Whilst these strategies have been clinically successful in patients with moderate-to-severe asthma (Piper et al., 2013), clinical trials in IBD patients have been inconclusive. IL-13 neutralisation by the human IgG4 monoclonal antibody tralokinumab increases the rate of clinical remission in patients with UC compared to controls (Danese et al., 2015), whereas anrukinzumab, a humanised IgG1 antibody, failed to alter clinical symptoms or induce remission (Reinisch et al., 2015).

IL-13 has recently been shown to directly activate sensory neurons which express IL-13Rα1 and IL-4Rα, and activation of neuronal IL-4Rα by IL-13 sensitizes sensory neurons to multiple other compounds (Oetjen et al., 2017). In colonic sensory neurons, transcript levels for both IL-13Rα1 and IL-4Rα are abundant and levels corelate with TRPV1 transcript levels, suggesting that IL-13 has the ability to interact with nociceptors innervating the colon (Hockley et al., 2019). Furthermore, transient receptor potential (TRP) channel-dependent IL-13 signalling has been linked to p38 MAPK and JAK activation (Oetjen et al., 2017), with increasing use of JAK inhibitors in the clinic to reduce visceral pain in GI disease (Danese et al., 2019). We therefore hypothesised that the primary agonist of the type II IL-4R, IL-13 sensitises colonic nociceptors to noxious mechanical distension and capsaicin application, the mechanism for which is dependent upon p38 MAPK and JAK signalling pathways.

## Materials and Methods

### Ethical approval

All animal experiments were conducted in compliance with the Animals (Scientific Procedures) Act 1986 Amendment Regulations 2012 under Project Licence P7EBFC1B1 granted to E. St. J. Smith by the Home Office with approval from the University of Cambridge Animal Welfare Ethical Review Body.

### Reagents

Stock concentrations of IL-13 (1 μM; H_2_O with 0.1% (w/v) bovine serum albumin) and capsaicin (1 mM; 100% ethanol) were purchased from ThermoFisher and Sigma-Aldrich respectively and dissolved as described. SB203580 (10 mM; DMSO), pyridone 6 (1 mM; DMSO) and ruxolitinib (1 mM; DMSO) were purchased from Tocris and stock concentrations made up as described. Nifedipine (100 mM; DMSO) and atropine (100 mM; 100% ethanol) were purchased from Sigma-Aldrich and dissolved as described. All drugs were diluted to working concentrations in extracellular solution (ECS) or Krebs buffer on the day of the experiment.

### Animals

Adult male C57BL/6J mice (8-16 weeks) were obtained from Charles River (Cambs, UK; RRID:IMSR_JAX:000664). Mice were conventionally housed in temperature-controlled rooms (21°C) with a 12-h light/dark cycle and provided with nesting material, a red plastic shelter and access to food and water ad libitum.

### Primary culture of mouse dorsal root ganglion neurons

DRG neurons were cultured as previously described (Barker et al., 2022). In brief, mice were humanely euthanised by exposure to a rising concentration of CO_2_, with confirmation by cervical dislocation. The spine was removed and bifurcated to allow isolation of DRG. Spinal DRG innervating the distal colon (T12-L5) were removed into ice-cold Leibovitz’s L-15 Medium, GlutaMAX™ Supplement (supplemented with 2.6% (v/v) NaHCO_3_). DRG were then incubated with 1 mg/ml collagenase (15 min) followed by 30 min trypsin (1 mg/ml) both with 6 mg/ml bovine serum albumin (BSA) in Leibovitz’s L-15 Medium, GlutaMAX™ Supplement (supplemented with 2.6% (v/v) NaHCO_3_). Following removal of the enzymes, DRG were resuspended in 2 ml Leibovitz’s L-15 Medium, GlutaMAX™ Supplement containing 10% (v/v) foetal bovine serum (FBS), 2.6% (v/v) NaHCO_3_, 1.5% (v/v) glucose and 300 units/ml penicillin and 0.3 mg/ml streptomycin (P/S). DRG were mechanically triturated using a 1 ml Gilson pipette and centrifuged (1000 rpm) for 30 s. The supernatant, containing dissociated DRG neurons, was collected in a separate tube and the pellet resuspended in 2 ml Leibovitz’s L-15 Medium, GlutaMAX™ Supplement containing 10% (v/v) FBS, 2.6% (v/v) NaHCO_3_, 1.5% (v/v) glucose and P/S. This process of mechanical trituration was repeated six times, after which the total supernatant was centrifuged (1000 rpm) for 5 min to pellet the dissociated DRG neurons. Cells were resuspended in 250 μl Leibovitz’s L-15 Medium, GlutaMAX™ Supplement containing 10% (v/v) FBS, 2.6% (v/v) NaHCO_3_, 1.5% (v/v) glucose and P/S and plated (50 μl per dish) onto 35 mm glass bottomed dishes coated with poly-D-lysine (MatTek, MA, USA) and laminin (Thermo Fisher: 23017015). Dishes were incubated for 3 hours (37°C, 5% CO_2_) to allow cell adhesion and subsequently flooded with Leibovitz’s L-15 Medium, GlutaMAX™ Supplement containing 10% (v/v) FBS, 2.6% (v/v) NaHCO_3_, 1.5% (v/v) glucose and P/S and incubated overnight. In experiments assessing the ability of pro-inflammatory cytokines to sensitise the capsaicin-induced Ca^2+^ response in DRG neurons, cultures were incubated overnight with TNFα (3 nM), IL-13 (30 nM) or phosphate-buffered saline (PBS) vehicle after flooding.

### Ca^2+^ imaging

Extracellular solution (ECS; in mM: 140 NaCl, 4 KCl, 1 MgCl_2_, 2 CaCl_2_, 4 glucose, 10 HEPES) was prepared and adjusted to pH 7.4 using NaOH and an osmolality of 290-310 mOsm with sucrose. Medium was aspirated from DRG neuronal cultures and cells were incubated (30 min) with 100 μl of 10 μM Ca^2+^ indicator Fluo-4-AM diluted in ECS (room temperature; shielded from light). For inhibitor studies requiring pre-incubation, 200 μl of drug was added for 10 min prior to imaging.

Dishes were mounted on the stage of an inverted microscope (Nikon Eclipse TE-2000S) and cells were visualised with brightfield illumination at 10 × magnification. To ensure a rapid exchange of solutions during protocols, the tip of a flexible perfusion onflow tube (AutoMate Scientific, CA, USA) attached to a six-channel, gravity-fed perfusion system (Warner Instruments, CT, USA) was placed beside the field of view. Initially, cells were superfused with ECS or drug in inhibitor studies to establish baseline.

Fluorescent images were captured with a CCD camera (Rolera Thunder, Qimaging, MC, Canada or Retiga Electro, Photometrics, AZ, USA) at 2.5 fps with 100 ms exposure. Fluo-4 was excited by a 470 nm light source (Cairn Research, Faversham, UK). Emission at 520 nm was recorded with μManager (Edelstein et al., 2014). All protocols began with a 10 s baseline of ECS before mediator superfusion, after which neurons were perfused with ECS for at least 20 s to remove the mediator. With multiple drug additions to the same dish, cells were allowed 4 min recovery between applications. Finally, cells were stimulated with 50 mM KCl for 10 s to determine cell viability, identify neuronal cells and allow normalisation of fluorescence. A fresh dish was used for each protocol and all solutions were diluted in ECS.

### Ca^2+^ imaging data analysis

Regions of interest were circled from a brightfield image and outlines overlaid onto fluorescent images using ImageJ (NIH, MA, USA). Pixel intensity was measured and analysed with custom-written scripts in RStudio (RStudio, MA, USA) to compute the proportion of neurons stimulated by each drug application and the corresponding magnitude of response. Briefly, the background fluorescence was subtracted from each region of interest, and fluorescence intensity (F) baseline corrected and normalised to the maximum fluorescence elicited during 50 mM KCl stimulation (F_pos_). Maximum KCl fluorescence was denoted as 1 F/F_pos_. Further analysis was confined to cells with a fluorescence increase ≥ 5 standard deviations above the mean baseline before 50 mM KCl application, as these were considered neurons. Using manual quality control, neurons were deemed responsive to drug application if a fluorescence increase of 0.1 F/F_pos_ was observed in response to drug perfusion. Responses to drug application were discounted if the increase in fluorescence began before or after the period of drug perfusion. The proportion of responsive neurons and magnitude of the fluorescence response was measured for each experiment, with peak responses calculated from averaging fluorescence values of individual neurons at each time point. Cell diameter was also measured with ImageJ software to allow size comparison of responsive neurons.

### Ex vivo whole-nerve electrophysiological recordings of colonic splanchnic afferents

LSN afferent preparations were conducted as previously described (Barker et al., 2022). Briefly, the distal colorectum (splenic flexure to rectum) and associated LSN (rostral to inferior mesenteric ganglia) were isolated from mice euthanised as described above. The colon was cannulated with fine thread (polyester, Gutermann) in a rectangular recording chamber with Sylgard base (Dow Corning, UK) and serosally superfused (7 ml/min; 32-34°C) and luminally perfused (200 μl/min) by a syringe pump (Harvard apparatus, MA) with carbogenated Krebs buffer solution (95% O_2_, 5% CO_2_). Krebs buffer was supplemented with 10 μM atropine and 10 μM nifedipine to prevent smooth muscle activity (Ness & Gebhart, 1988a).

Borosilicate suction electrodes were used to record the multi-unit activity of LSN bundles. Signals were amplified (gain 5 kHz), band pass filtered (100–1300 Hz; Neurolog, Digitimer Ltd, UK), and digitally filtered for 50 Hz noise (Humbug, Quest Scientific, Canada). Analogue signals were digitized at 20 kHz (Micro1401; Cambridge Electronic Design, UK) and signals were visualised with Spike2 software (Cambridge Electronic Design, UK).

### Electrophysiology Protocols

Following dissection, preparations underwent a minimum 30 min stabilisation period to ensure baseline firing was consistent. Repeated ramp distensions (0-80 mmHg) were performed on the cannulated distal colon by occluding the luminal perfusion out-flow and increasing intraluminal pressure gradually over approximately 220 s. When the maximum 80 mmHg pressure was achieved, the luminal outflow was re-opened and pressure returned to baseline. Distension pressures > 30mmHg evoke noxious pain behaviours in mice and humans (Hughes, Brierley, Martin, et al., 2009; Ness & Gebhart, 1988b).

Each protocol consisted of five ramp distensions, separated by 15 min to allow baseline stabilisation, after which the colon was superfused with 1 μM capsaicin (20 ml). Between ramps 3 and 4, mediator (100 nM TNFα or 30 nM IL-13) or vehicle (buffer) was applied by luminal perfusion for 15 min. In protocols examining the effects of pathway inhibition, preparations were pre-treated with inhibitor (10 μM SB203580 or 1 μM Pyridone 6) or vehicle (0.01% DMSO) for 15 min before and throughout mediator perfusion.

### Electrophysiological Data Analysis

Nerve discharge was determined by quantifying the number of spikes passing through a manually determined threshold, twice the level of background noise (typically 60-80 μV) and binned to determine the average firing frequency over 10 s. Baseline firing was established by the average activity 180 s prior to distension or drug perfusion. Changes to neuronal firing rates were calculated by subtracting baseline activity from increases in nerve activity in response to ramp distension or perfusion with capsaicin. Peak firing to noxious mechanical distension and capsaicin application was defined respectively as the highest neuronal activity during the fifth ramp distension and during the 10 min post capsaicin application. Changes to neuronal activity during ramp distension were measured at each 5 mmHg increase in pressure and used to visualise ramp profiles. Response profiles to capsaicin application were plotted from binned data averaged from 30 s increments. The area under the curve (AUC) was calculated for the duration of each ramp distension (0-80 mmHg) and for the 10 min post initial capsaicin application following the generation of response profiles using GraphPad Prism 9 software (GraphPad Software, CA).

### Statistical Analysis

All data sets were normality tested with a Shapiro-Wilk test and analysed using the appropriate statistical tests. The level of significance was set at p ≤ 0.05. All data are displayed as means ± standard deviation (SD). For Ca^2+^ imaging analysis, n represents the total number of dishes and N represents the total number of mice from which they were cultured. On average, ∼180 neurons were imaged per dish.

## Results

### IL-13 increases [Ca^2+^]_i_ in DRG sensory neurons co-sensitive to capsaicin

Previous studies investigating the role of IL-13 in itch sensory pathways have shown its ability to directly activate DRG sensory neurons (Oetjen et al., 2017). To corroborate these findings and the extent to which IL-13 stimulated DRG sensory neurons, 30 nM IL-13 was applied to neurons and the change in [Ca^2+^]_i_ measured using Fluo4. As expected, IL-13 increased [Ca^2+^]_i_ as demonstrated by an increase in Fluo4 fluorescence (Figure 1A). Of the total neurons assessed, IL-13 activated 24.8 ± 12.1%, significantly fewer than those that were IL-13-insensitive (p = 0.0001, unpaired t-test; n = 5 N = 5; Figure 1B). Comparison of cell size also revealed that IL-13 stimulated a population of smaller diameter neurons than those that were non-responsive (p < 0.0001, Kolmogorov-Smirnov test; n = 644 neurons; Figure 1C), suggesting that IL-13 was preferentially stimulating nociceptors. To investigate this further, IL-13-sensitive neurons were evaluated for their co-sensitivity to 1 μM capsaicin (Figure 1Di). Three populations of responsive neurons were identified, including those that were co-sensitive to capsaicin and IL-13 (Figure 41Dii). In total, 27.3 ± 11.2% of neurons were activated by IL-13, 89.1% of which were also sensitive to capsaicin (Figure 1E). Overall, 40.8 ± 9.26% of neurons were sensitive to capsaicin, but only 59.7% of these were activated by IL-13, suggesting that IL-13 does not stimulate all nociceptors. This is corroborated by cell diameter data showing that neurons responsive to capsaicin alone (p < 0.0001) and those with overlapping sensitivity to capsaicin and IL-13 were smaller than neurons unresponsive to both IL-13 and capsaicin (p < 0.0001, Kruskal-Wallis test with Dunn’s multiple comparison; n = 488 neurons; Figure 1F).

**Figure 1:**
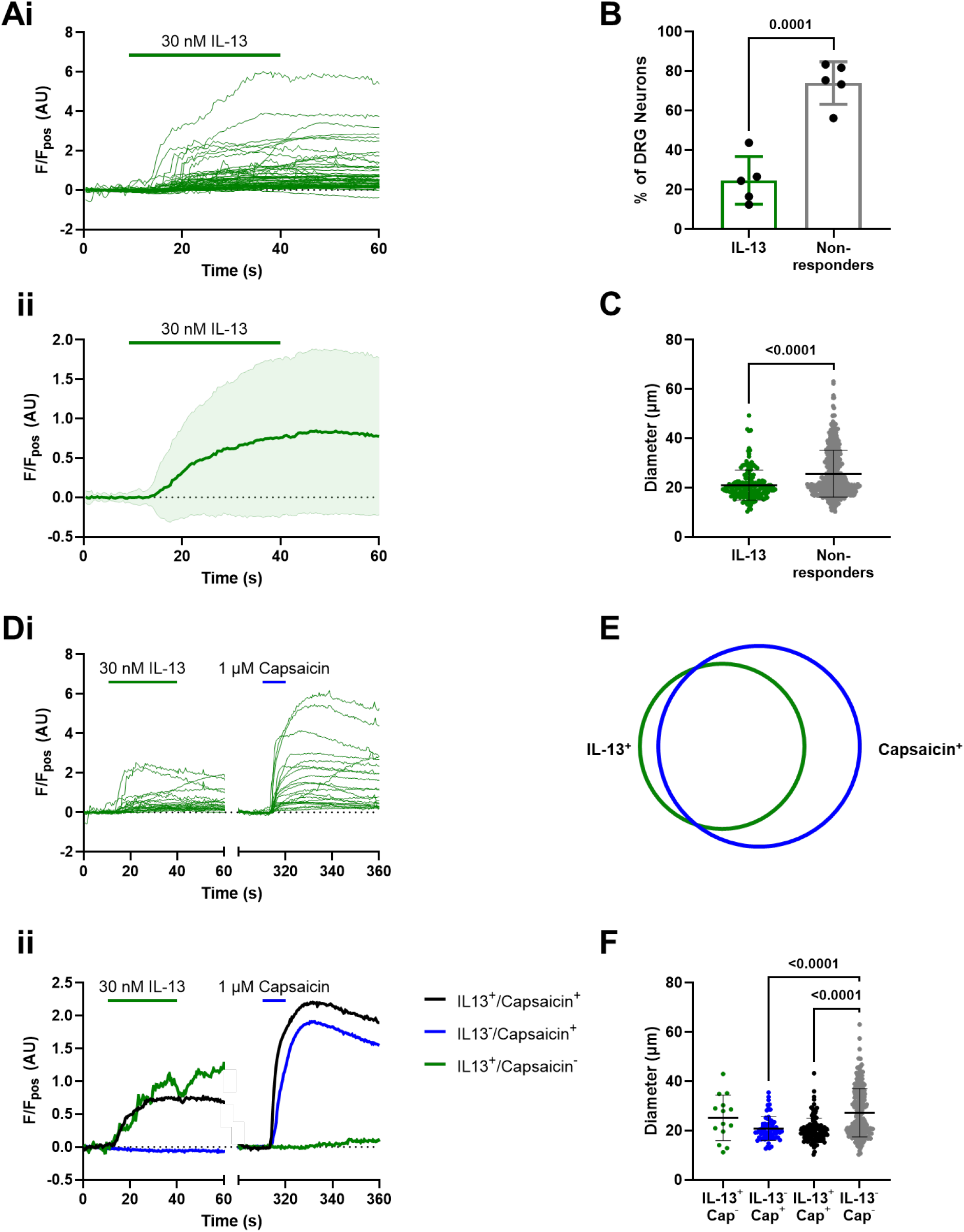
IL-13 increases [Ca^2+^]_i_ in capsaicin-sensitive DRG neurons. (A) Representative traces from individual (i) and averaged (ii) neuronal responses to 30 nM IL-13 (n = 63 neurons). (B) The percentage of DRG neurons stimulated by IL-13 was significantly lower than non-responsive neurons (p = 0.0001, unpaired t-test; n = 5 N = 5). (C) IL-13-sensitive neurons were smaller in diameter than non-responders (p < 0.0001, Kolmogorov-Smirnov test; n = 644 neurons). (D) Example traces from (i) individual neurons responsive to both IL-13 and 1 μM capsaicin and (ii) response profiles of responder populations. (E) Representative Venn diagram of overlapping responses in neurons stimulated with IL-13 and capsaicin (n = 5, N = 5). (F) Size distribution of IL-13-and/or capsaicin-sensitive neurons. IL/13/capsaicin co-sensitive and capsaicin-sensitive neurons were smaller in diameter than non-responders (p < 0.0001, p < 0.0001, Kruskal-Wallis test with Dunn’s multiple comparison; n = 488 neurons).

### IL-13-mediated Ca^2+^ responses are dependent on JAK and p38 MAPK signalling

Activation of IL-13 receptor components IL-4Rα or IL-13Rα initiates transphosphorylation of JAK proteinsand initiates signal transduction in a range of cell types, including immune cells (Kelly-Welch et al., 2003; Malabarba et al., 1996; Miyazaki et al., 1994; Roy & Cathcart, 1998; Welham et al., 1995). In sensory neurons, activation of IL-4Rα by IL-4 is dependent upon JAK1 signalling (Oetjen et al., 2017). To determine whether JAK signalling is also involved in the activation of DRG sensory neurons by IL-13, cultures were incubated with the pan-JAK inhibitor, pyridone 6 (1 μM). JAK inhibition decreased the proportion of IL-13-responsive neurons from 20.8 ± 5.62% to 7.58 ± 4.17% (p = 0.003, one-way ANOVA with Holm-Šídák’s multiple comparisons test; n = 5; Figure 2A). Following this, neurons were incubated with 1 μM ruxolitinib, a selective JAK1/2 inhibitor, to help establish more specifically which JAK signalling proteins were involved in IL-13-mediated increases in [Ca^2+^]_i_. Ruxolitinib reduced the proportion of IL-13 responders, with 7.20 ± 6.60% of neurons still activated (p = 0.002, one-way ANOVA with Holm-Šídák’s multiple comparisons test; n = 5; Figure 2A), which was equivalent to pyridone 6 treatment (p > 0.999, one-way ANOVA with Holm-Šídák’s multiple comparisons test; n = 5), suggesting that JAK1 and JAK2 are involved in DRG sensory neuron activation by IL-13.

**Figure 2:**
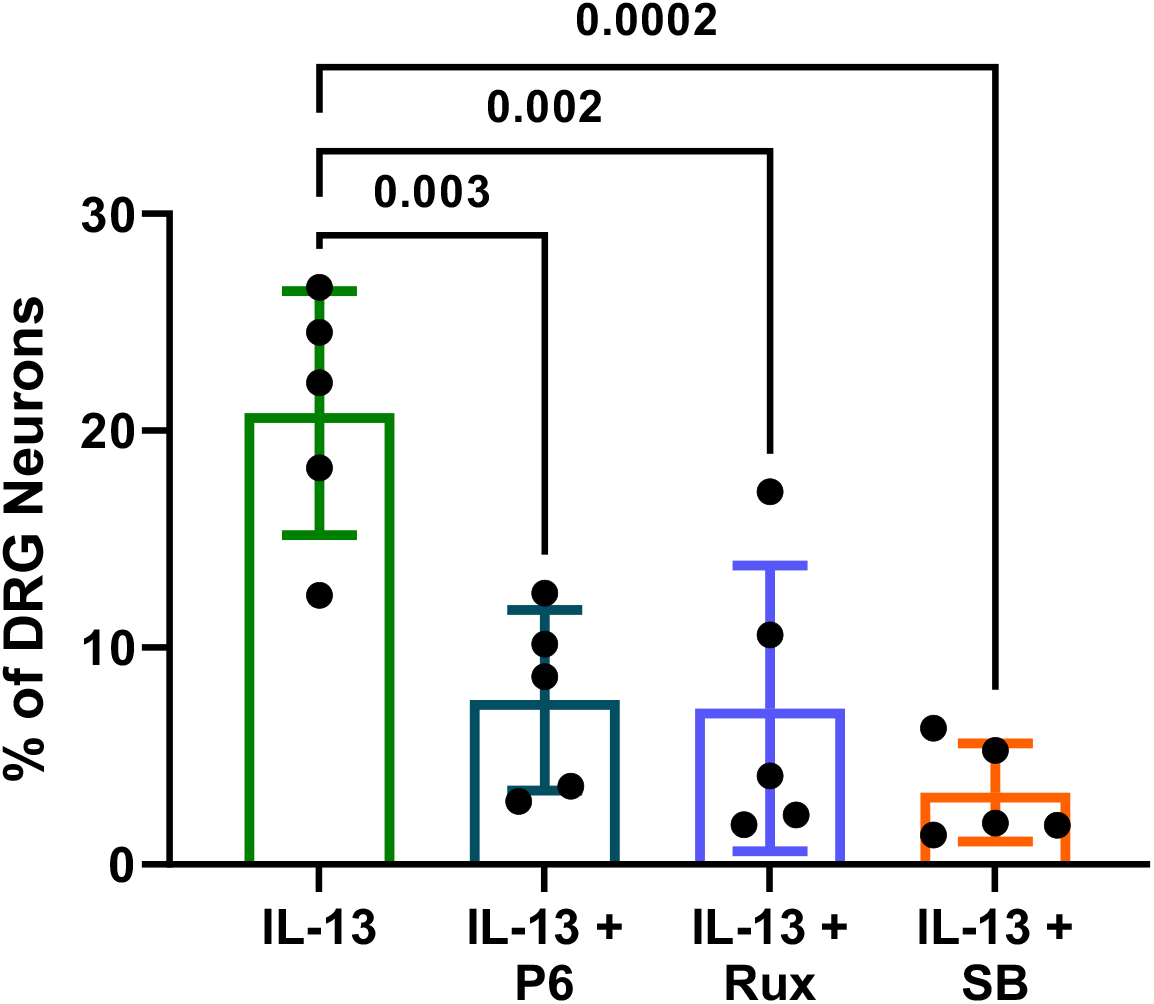
Activation of DRG sensory neurons by IL-13 is JAK1/2- and p38 MAPK-dependent. The proportion of IL-13-sensitive neurons was significantly reduced by pan-JAK inhibition with pyridone 6 (P6) (p = 0.003, one-way ANOVA with Holm-Šídák’s multiple comparisons test; n = 5), JAK1/2 inhibition with ruxolitinib (Rux) (p = 0.002) and p38 MAPK inhibition with SB203580 (SB) (p = 0.0002).

Aside from JAK signalling, p38 MAPK phosphorylation/activation has also been observed in immune cells following exposure to IL-13 and, in mast cells, IL-13 production is p38 MAPK-dependent, forming a feedback loop during inflammation (Petrova et al., 2020; Xu et al., 2003). To investigate if there was a role for p38 MAPK in IL-13 signalling in DRG sensory neurons, the p38 MAPK inhibitor SB203580 was added to culture dishes. In line with those observations in immune cells, p38 MAPK inhibition reduced the proportion of IL-13 responsive neurons by 85% (p = 0.0002, one-way ANOVA with Holm-Šídák’s multiple comparisons test; n = 5; Figure 2A).

### IL-13 sensitises colonic afferent responses to noxious ramp distension

Stable responses of the LSN to colonic distension were achieved after 2-3 distensions. Thus ramp 3 was used as a control ramp in each protocol, after which tissues were perfused with 100 nM IL-13 for 15 min and the effects on LSN response to mechanical distension observed (Figure 3A). In vehicle-treated tissues, no significant difference was observed in the AUC between distension 3 and distension 5 responses (p = 0.708, two-way ANOVA with Holm-Šídák’s multiple comparisons test; N = 8; Figure 3B). Similarly, in tissues perfused with IL-13, responses to distension were unchanged between the third and fifth ramp distensions (p = 0.708, two-way ANOVA with Holm-Šídák’s multiple comparisons test; N = 8). Furthermore, between treatment groups, distension 3 and distension 5 responses were no different (p = 0.708, p = 0.157 respectively, two-way ANOVA with Holm-Šídák’s multiple comparisons test; N = 8). Despite this, a significant increase in nerve activity in response to ramp distension occurred across distending pressures (50-80 mmHg) in IL-13-treated tissues (p = 0.003, multiple unpaired t-tests, N = 8; Figure 3C), with the greatest difference in nerve firing observed at 80 mmHg (vehicle: 29.7 ± 4.62 spikes s^−1^ vs IL-13: 40.5 ± 6.97 spikes s^−1^; p = 0.003, unpaired t-test, N = 8). Furthermore, IL-13 elevated peak afferent firing in response to noxious ramp distension by over 11 spikes s^−1^ compared to vehicle, with average peak activity of 41.2 ± 7.26 spikes s^−1^ (p = 0.002, unpaired t-test, N = 8; Figure 3D). Unlike in DRG neurons where direct activation by IL-13 was observed, IL-13 failed to alter basal LSN activity *ex vivo* (p = 0.101, unpaired t-test; N = 8; Figure 3E), and a similar rate of spontaneous firing was maintained between treatment groups (vehicle: 15.6 ± 5.38 spikes s^−1^ vs IL-13: 11.7 ± 3.39 spikes s^−1^). Therefore, in LSN colonic afferents, IL-13 sensitises responses to noxious mechanical stimuli, but does not cause direct activation, unlike in DRG neurons where direct activation is seen, though mechanosensitivity was not measured. Compliance of ramp distension was measured in each protocol and, between treatment groups, ramp distensions 3 (vehicle: 0.414 ± 0.156 ml vs IL-13: 0.566 ± 0.285 ml) and 5 (vehicle: 0.415 ± 0.154 ml vs IL-13: 0.483 ± 0.131 ml) required the same volume of fluid to increase intraluminal pressure to 80 mmHg (p = 0.544, p = 0.862 respectively, two-way ANOVA with Holm-Šídák’s multiple comparisons test; N = 8; Figure 3F). Moreover, no significant difference in compliance was measured between distension 3 and 5 within treatment groups (control: p = 0.987, IL-13: p = 0.862, two-way ANOVA with Holm-Šídák’s multiple comparisons test; N = 8).

**Figure 3:**
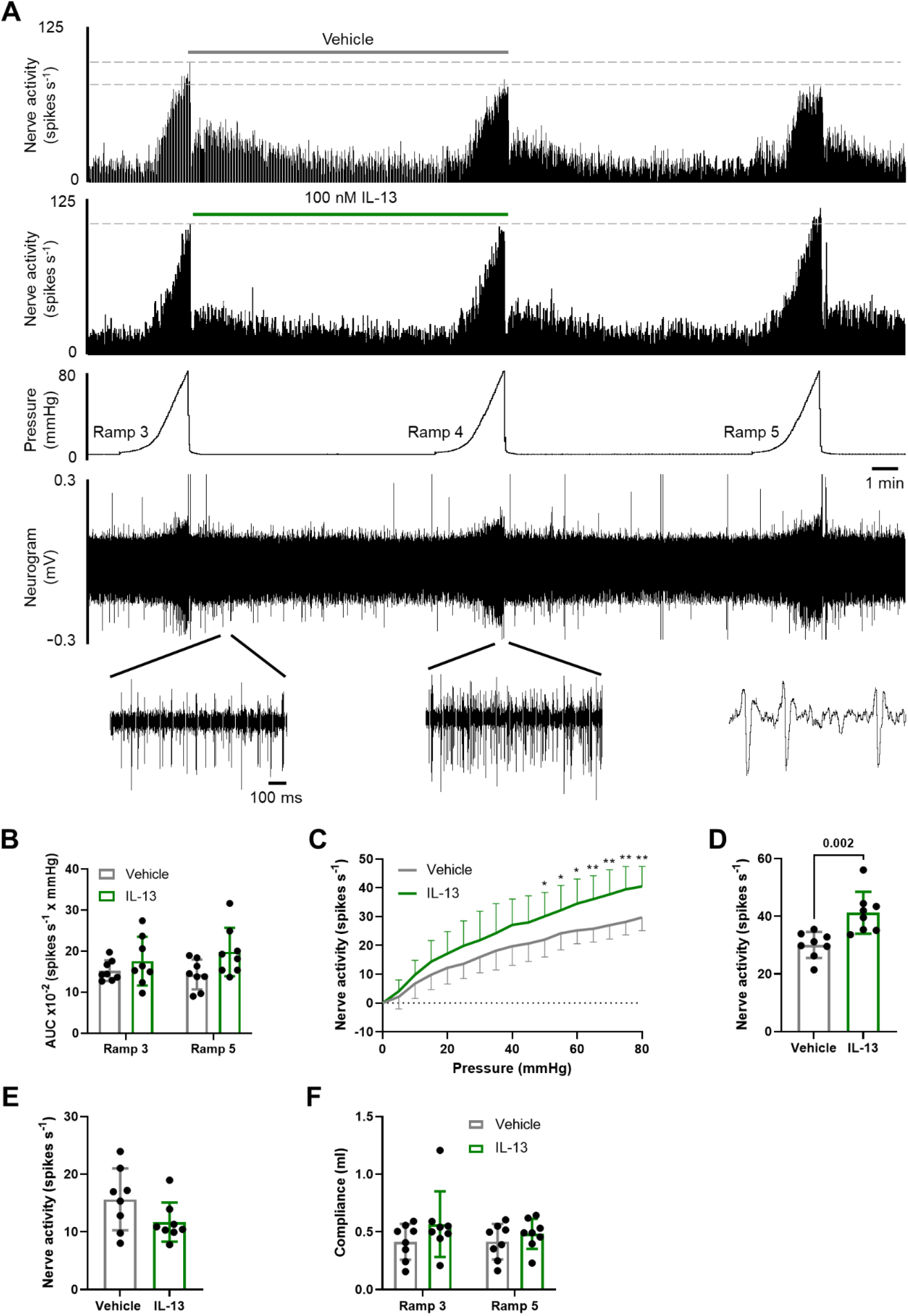
IL-13 evokes colonic afferent mechanosensitivity to noxious ramp distension. (A) Example rate histograms and neurogram of LSN activity with accompanying pressure trace showing sequential (x3) ramp distensions (0-80 mmHg) from vehicle- and IL-13 (100 nM)-treated preparations, demonstrating the sensitisation of colonic afferent responses to noxious distension following IL-13 perfusion. (B) Area under the curve (AUC) of colonic afferent responses to distension was unchanged in IL-13-treated tissues compared with vehicle (ramp 5) (p = 0.708, two-way ANOVA with Holm-Šídák’s multiple comparisons test; N = 8). (C) IL-13 increased afferent firing across a range of noxious distending pressures (> 50mmHg, ramp 5) compared to vehicle (* p < 0.05, ** p < 0.01, multiple t-tests; N = 8). (D) Peak afferent activity was significantly increased in tissues following IL-13 pre-treatment (p = 0.002, unpaired t-test; N = 8). (E) IL-13 did not directly alter spontaneous afferent activity (p = 0.101, unpaired t-test; N = 8). (F) Compliance was unchanged between sequential ramp distensions within treatment groups (vehicle: p = 0.987, IL-13: p = 0.862, two-way ANOVA with Holm-Šídák’s multiple comparisons test; N = 8) and no differences between treatment groups were observed (ramp 3: p = 0.544, ramp 5: p = 0.862, two-way ANOVA with Holm-Šídák’s multiple comparisons test; N = 8).

### IL-13 does not sensitise colonic afferent response to noxious capsaicin application

To determine whether, as well as sensitising colonic afferents to mechanical stimuli, IL-13 also sensitised TRPV1-mediated LSN activity, preparations were perfused with 1 μM capsaicin (20 ml). As seen previously, capsaicin produced a direct increase in nerve activity in vehicle-treated tissues (Figure 4A). A similar capsaicin response was observed in preparations treated with 100 nM IL-13, represented by equivalent response AUCs (p = 0.818, unpaired t-test; N = 8; Figure 4B). IL-13 did not sensitise afferent activity in the 10 min post capsaicin application (p = 0.979, two-way ANOVA with Holm-Šídák’s multiple comparisons test; N = 8; Figure 4C) and peak afferent firing remained unchanged between treatment groups (vehicle: 32.4 ± 3.44 spikes s^−1^, IL-13: 34.3 ± 7.92 spikes s^−1^; p = 0.534, unpaired t-test; N = 8; Figure 4D). Furthermore, tissues treated with IL-13 (411 ± 87.1 s) and vehicle (392 ± 83.9 s) required the same amount of time to return to basal nerve activity post capsaicin application (p = 0.649, unpaired t-test; N = 8; Figure 4E). Together, these data show that, whilst IL-13 sensitised visceral afferents to noxious mechanical distension, IL-13 does not sensitise TRPV1 in colonic afferents.

**Figure 4:**
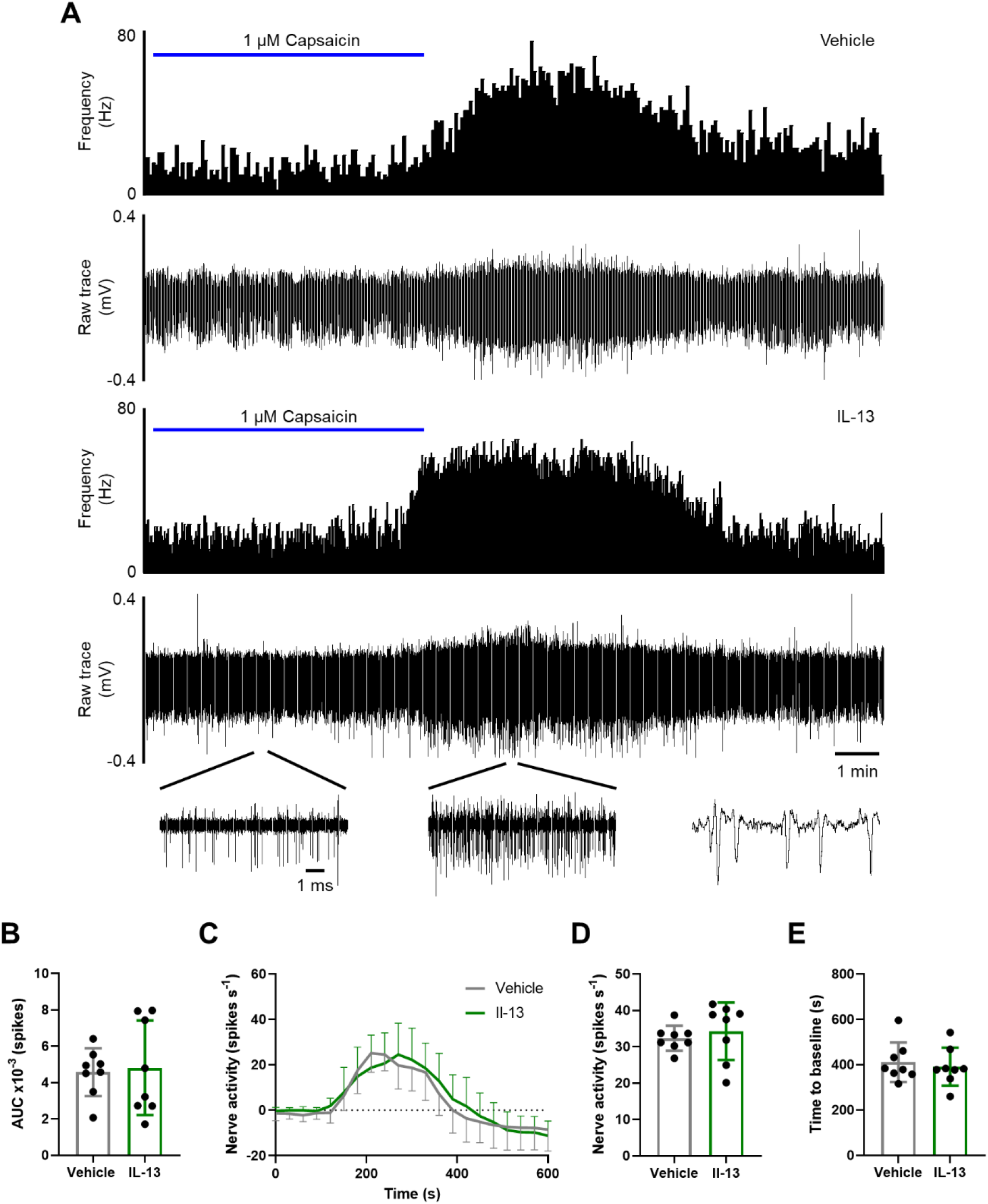
IL-13 does not sensitise colonic afferent responses to capsaicin. (A) Example rate histograms and accompanying neurograms of LSN activity, illustrating similar afferent responses to 1 μM capsaicin following incubation with IL-13 (100 nM) compared with vehicle pre-treatment. (B) IL-13 did not alter response AUC 0-10 min post capsaicin application (p = 0.818, unpaired t-test; N = 8). (C) Afferent firing was unchanged in the 10 min post capsaicin application (p = 0.979, two-way ANOVA with Holm-Šídák’s multiple comparisons test; N = 8). (D) Peak afferent firing remained constant between vehicle- and IL-13-treated tissues (p = 0.534, unpaired t-test; N = 8). (E) The time taken for nerve activity to return to baseline firing following capsaicin application was unchanged by IL-13 incubation (p = 0.649, unpaired t-test; N = 8).

### JAK signalling mediates IL-13-induced hypersensitivity to noxious mechanical distension

Following on from experiments in DRG neurons showing IL-13-mediated activation was dependent on JAK signalling, we next explored the role of JAK in colonic afferent sensitisation to mechanical distension. The pan-JAK inhibitor pyridone 6 (1 μM) was applied to LSN preparations in the presence of either vehicle or IL-13. Pyridone 6 decreased the AUC of afferent responses to noxious ramp distension following pre-incubation with 100 nM IL-13 (p = 0.046, one-way ANOVA with Holm-Šídák’s multiple comparisons test; n = 6-8; Figure 5A). Preparations perfused with vehicle and pyridone 6 also showed significantly reduced responses to ramp distension compared to IL-13 (p = 0.011, one-way ANOVA with Holm-Šídák’s multiple comparisons test; n = 6-8). It was further observed that pan-JAK inhibition attenuated IL-13-induced sensitisation across noxious distending pressures (p = 0.012, multiple unpaired t-tests; N = 6-8; Figure 5B). Additionally, pyridone 6 alone significantly reduced ramp distension responses compared to IL-13-treated tissues (p = 0.003, multiple unpaired t-tests; N = 6-8), with responses no different from those seen in vehicle-treated tissues across distending pressures (p = 0.069, multiple unpaired t-tests; N = 6-8). Pyridone 6 incubation also significantly decreased peak afferent firing to distension with an average activity of 29.2 ± 10.1 spikes s^−1^ in the presence of IL-13 (p = 0.038, one-way ANOVA with Holm-Šídák’s multiple comparisons test; N = 6-8; Figure 5C) and peak afferent firing was also reduced in pyridone 6 controls (24.2 ± 9.94 spikes s^−1^; p = 0.008, one-way ANOVA with Holm-Šídák’s multiple comparisons test; N = 6-8). Spontaneous firing rates in IL-13-treated tissues were unaffected by pyridone 6 (p > 0.999, Kruskal-Wallis test with Dunn’s multiple comparisons test; N = 6-8; Figure 5D), as demonstrated by baseline activity being similar across all treatment groups (pyridone 6: 9.22 ± 4.71 spikes s^−1^, pyridone 6 + IL-13: 13.4 ± 8.10 spikes s^−1^; pyridone 6 vs IL-13: p > 0.999, pyridone 6 vs pyridone 6 + IL-13: p > 0.999, Kruskal-Wallis test with Dunn’s multiple comparisons test; N = 6-8). Compliance was also assessed to ensure consistency between distensions (Figure 5E). Compliance of distension 3 was not significantly different from distension 5 in either pyridone 6- (ramp 3: 0.547 ± 0.095 ml, ramp 5: 0.696 ± 0.346 ml; p = 0.972) or pyridone 6 + IL-13-treated preparations (ramp 3: 0.571 ± 0. 0.178 ml vs ramp 5: 0.505 ± 0.154 ml, p = 0.999, two-way ANOVA with Holm-Šídák’s multiple comparisons test; N = 6-8). Similar volumes were also required for distension 3 (pyridone 6 vs pyridone 6 + IL-13: p > 0.999; pyridone 6 vs IL-13: p > 0.999; pyridone 6 + IL-13 vs IL-13: p > 0.999) and distension 5 between treatment groups (pyridone 6 vs pyridone 6 + IL-13: p = 0.834; pyridone 6 vs IL-13: p = 0.688; pyridone 6 + IL-13 vs IL-13: p > 0.999, two-way ANOVA with Holm-Šídák’s multiple comparisons test; N = 6-8).

**Figure 5:**
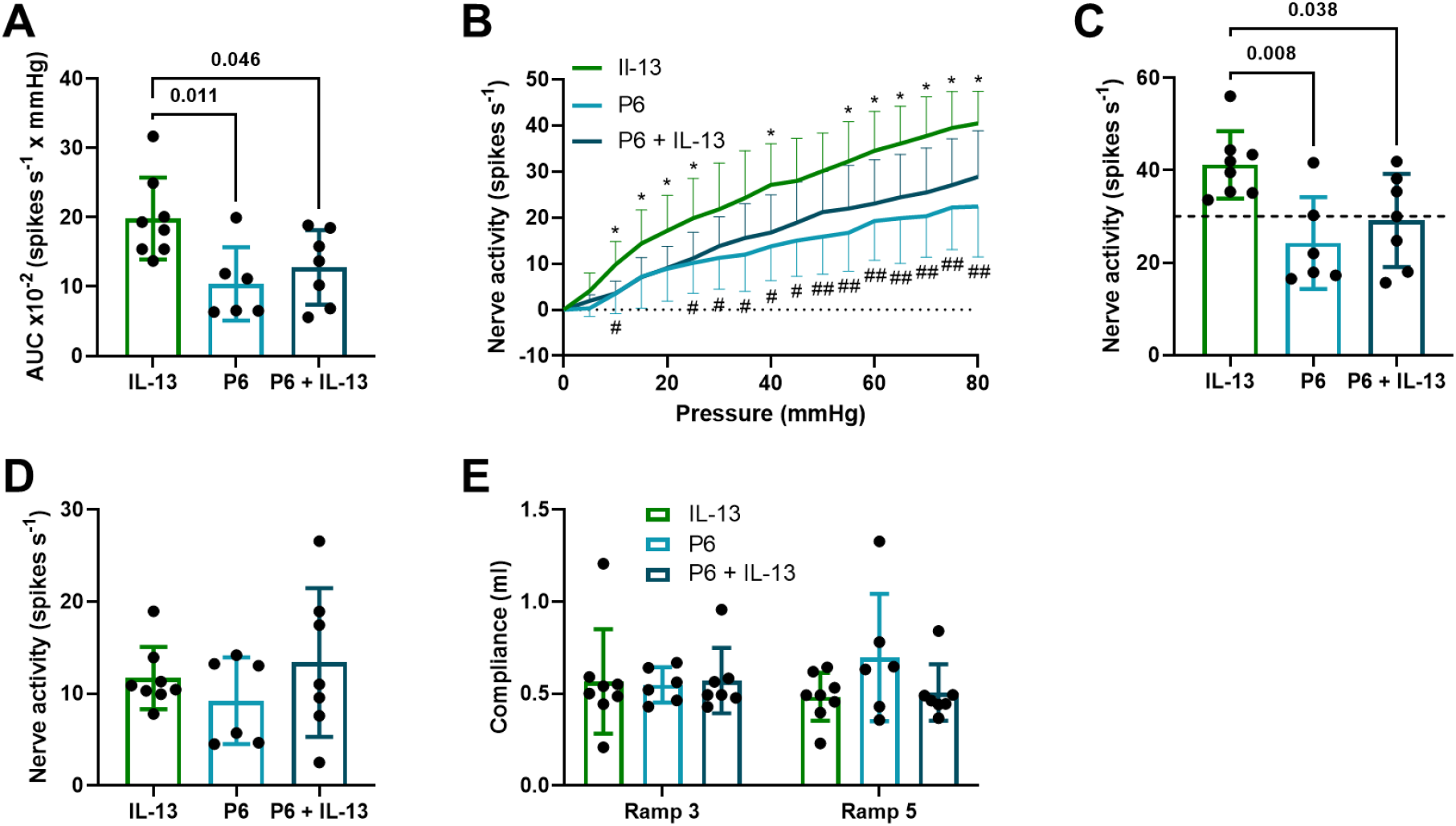
IL-13-mediated sensitisation of colonic afferents to mechanical ramp distension is JAK mediated. (A) Pan-JAK inhibition with 1 μM pyridone 6 (P6) reduced afferent responses to noxious distension compared to IL-13-treated tissues (p = 0.046, one-way ANOVA with Holm-Šídák’s multiple comparisons test; N = 6-8). (B) JAK inhibition reduced afferent responses across distending pressures (> 10 mmHg) (IL-13 vs pyridone 6 + IL-13: * p < 0.05; IL-13 vs pyridone 6: # p < 0.05, ## p < 0.01, multiple t-tests; N = 6-8). (C) Pyridone 6 significantly reduced the increase in peak nerve activity in response to ramp distension following IL-13 incubation (p = 0.038, one-way ANOVA with Holm-Šídák’s multiple comparisons test; N = 6-8). Dashed line shows the mean nerve activity during vehicle application (see Figure 3). (D) Spontaneous afferent activity was unaffected by pyridone 6 (p > 0.999, Kruskal-Wallis test with Dunn’s multiple comparisons test; N = 6-8). (E) Compliance was unchanged between ramp 3 and ramp 5 within treatment groups (pyridone 6: p = 0.972, pyridone 6 + IL-13: p = 0.999, two-way ANOVA with Holm-Šídák’s multiple comparisons test; N = 7-8) and no difference was found in distension 3 responses (pyridone 6 vs pyridone 6 + IL-13: p > 0.999; pyridone 6 vs IL-13: p > 0.999; pyridone 6 + IL-13 vs IL-13: p > 0.999) or distension 5 responses between treatment groups (pyridone 6 vs pyridone 6 + IL-13: p = 0.834; pyridone 6 vs IL-13: p = 0.688; pyridone 6 + IL-13 vs IL-13: p > 0.999, two-way ANOVA with Holm-Šídák’s multiple comparisons test; N = 6-8).

### IL-13-mediated sensitivity to noxious mechanical distension is p38 MAPK-dependent

Considering that IL-13 activation was blocked by p38 MAPK inhibition in DRG neurons, the functional effect of SB203580 on LSN afferent response to noxious mechanical distension was assessed in *ex vivo* intact tubular colonic preparations. Firstly, as with JAK inhibition, no sensitisation of distension responses was seen following IL-13 (100 nM) incubation with the p38 MAPK inhibitor SB203580 (10 μM; p = 0.002, one-way ANOVA with Holm-Šídák’s multiple comparisons test; n = 6-8; Figure 6A) or in SB203580 controls (p = 0.003, one-way ANOVA with Holm-Šídák’s multiple comparisons test; n = 6-8). Furthermore, no difference in AUC was observed between SB203580-treated groups (p = 0.646, one-way ANOVA with Holm-Šídák’s multiple comparisons test; n = 6-8). However, inhibition of p38 MAPK attenuated IL-13-induced sensitisation across all distending pressures (5-80 mmHg; p = 0.002; multiple unpaired t-tests; N = 6-8; Figure 6B) and distension response activity was significantly reduced across all pressures (5-80 mmHg) in SB203580 controls compared to IL-13-treated tissues (p = 0.004, multiple unpaired t-tests; N = 6-8). Moreover, peak afferent firing in response to distension with IL-13 was attenuated in SB203580-treated tissues (28.9 ± 4.88 spikes s^−1^; p = 0.008, Kruskal-Wallis test with Dunn’s multiple comparisons test; N = 6-8; Figure 6C). SB203580 controls also demonstrated reduced peak afferent firing to ramp distension (29.5 ± 3.97 spikes s^−1^; p = 0.040, Kruskal-Wallis test with Dunn’s multiple comparisons test; N = 6-8) and SB203580-treated groups displayed equivalent peak responses (p > 0.999, Kruskal-Wallis test with Dunn’s multiple comparisons test; N = 6-8). Spontaneous nerve activity upon IL-13 addition remained unchanged in the presence of SB203580 (12.5 spikes s^−1^; p = 0.993, one-way ANOVA with Holm-Šídák’s multiple comparisons test; N = 6-8; Figure 6D). SB203580 controls also displayed similar basal firing (11.9 ± 3.98 spikes s^−1^) to IL-13 (p = 0.993) and SB203580 + IL-13 protocols (p = 0.993, one-way ANOVA with Holm-Šídák’s multiple comparisons test; N = 6-8). Ramp distension compliance was unchanged between distension 3 and 5 within SB203580 (ramp 3: 0.571 ± 0.112 ml, ramp 5: 0.570 ± 0.092 ml; p > 0.999) and SB203580 + IL-13 tissues (ramp 3: 0.612 ± 0.113 ml vs ramp 5: 0.568 ± 0.102 ml, p > 0.999, two-way ANOVA with Holm-Šídák’s multiple comparisons test; N = 6-8; Figure 6E). Compliance was also no different between third distensions (SB203580 vs SB203580 + IL-13: p > 0.999; SB203580 vs IL-13: p > 0.999; SB203580 + IL-13 vs IL-13: p > 0.999) and fifth distensions (SB203580 vs SB203580 + IL-13: p > 0.999; SB203580 vs IL-13: p = 0.995; SB203580 + IL-13 vs IL-13: p = 0.995, two-way ANOVA with Holm-Šídák’s multiple comparisons test; N = 6-8) across treatment groups.

**Figure 6:**
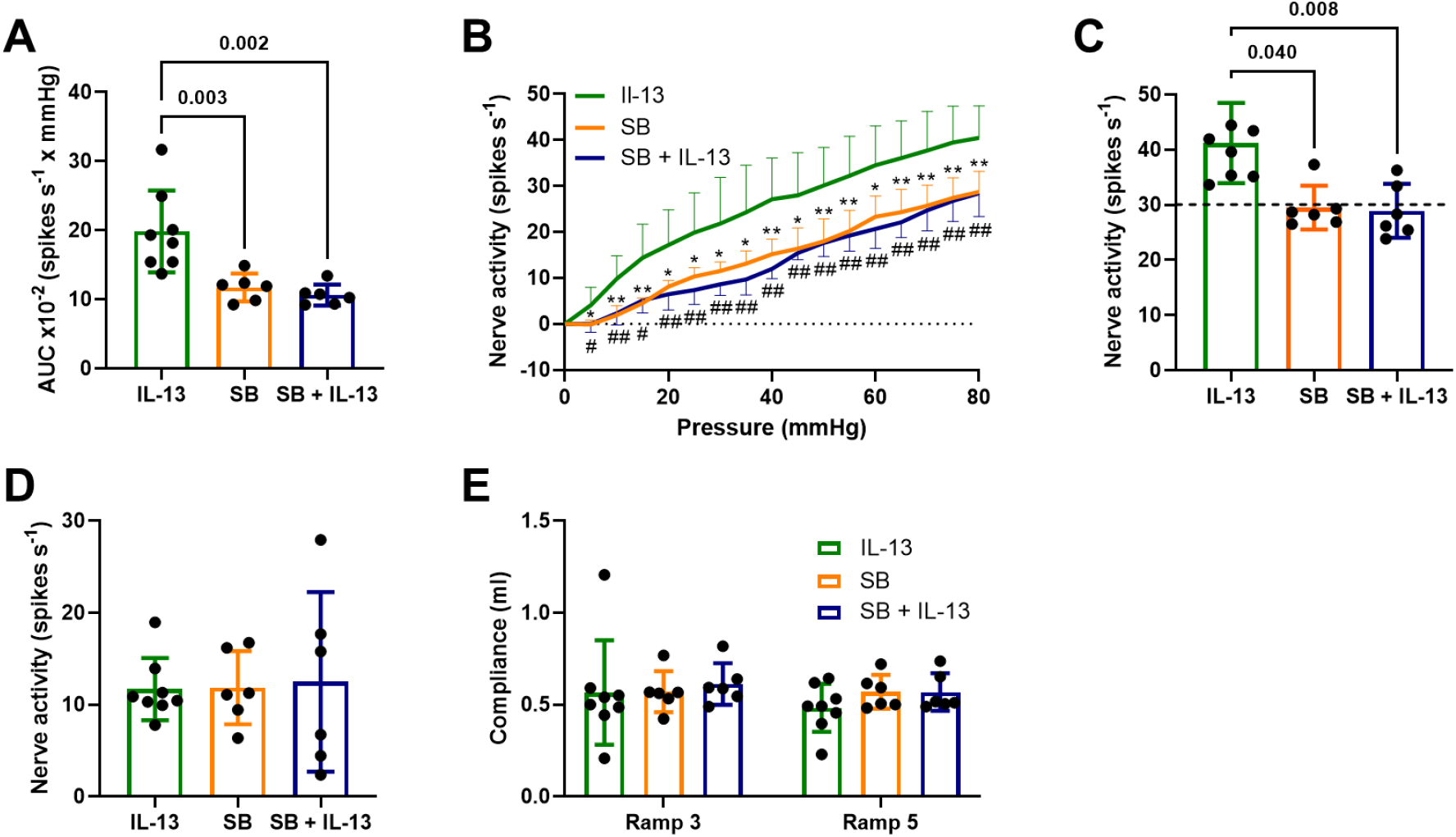
IL-13-induced colonic afferent sensitisation to mechanical ramp distension is dependent on p38 MAPK. (A) IL-13 (100 nM) failed to sensitise afferent responses to noxious distension in 10 μM SB203580 (SB)-treated tissues (p = 0.002, one-way ANOVA with Holm-Šídák’s multiple comparisons test; n = 6-8). (B) p38 MAPK inhibition reduced afferent responses across all distending pressures (> 5 mmHg) (IL-13 vs SB + IL-13: # p < 0.05, ## p < 0.01; IL-13 vs SB: * p < 0.05, ** p < 0.01, multiple t-tests; N = 6-8). (C) Peak nerve activity in response to noxious ramp distension was significantly reduced by SB203580 (p = 0.008, Kruskal-Wallis test with Dunn’s multiple comparisons test; N = 6-8). Dashed line shows the mean nerve activity during vehicle application (see Figure 3). (D) IL-13 had no direct effect on spontaneous afferent activity in tissues treated with SB203580 (p = 0.993, one-way ANOVA with Holm-Šídák’s multiple comparisons test; N = 6-8). (E) Compliance was unchanged between ramp 3 and ramp 5 within treatment groups (SB203580: p > 0.999, SB203580 + IL-13: p > 0.999, two-way ANOVA with Holm-Šídák’s multiple comparisons test; N = 6-8) and no difference was found between ramp 3 (SB203580 vs SB203580 + IL-13: p > 0.999; SB203580 vs IL-13: p > 0.999; SB203580 + IL-13 vs IL-13: p > 0.999) and ramp 5 responses (SB203580 vs SB203580 + IL-13: p > 0.999; SB203580 vs IL-13: p = 0.995; SB203580 + IL-13 vs IL-13: p = 0.995, two-way ANOVA with Holm-Šídák’s multiple comparisons test; N = 6-8) between treatment groups.

## Discussion

IL-13 is a pro-inflammatory cytokine associated with a number of inflammatory conditions, such as asthma (Doran et al., 2017), atopic dermatitis (Oetjen et al., 2017) and IBD (Vainer et al., 2000). With regard to pain, IL-13 has been observed to both activate and sensitise sensory neurons (Oetjen et al., 2017). As seen with TNFα, studies have shown that IL-13-mediated neuronal activation is dependent on extracellular Ca^2+^ entry via TRP channels, including TRPV1 (Oetjen et al., 2017). Despite this, no data exists to implicate IL-13 signalling in colonic afferent sensitisation, leaving little knowledge of IL-13 involvement in hypersensitivity and visceral pain in GI disease. We therefore investigated and found a pro-nociceptive role of IL-13 in colonic afferents, which was dependent upon p38 MAPK and JAK signalling in the sensitisation of responses to noxious mechanical distension. Overall, these data present IL-13 as a potential target for treating pain originating in the colon and highlight the possible benefit of targeting downstream p38 MAPK and JAK signalling pathways to alleviate pain.

Firstly, we observed an IL-13-evoked rise in [Ca^2+^]_i_ in DRG sensory neurons, in line with previous studies (Oetjen et al., 2017), the majority of which were small in diameter, thus likely nociceptors. We found that IL-13 activated a greater proportion of DRG neurons compared to previous studies, likely the result of differences in culture conditions, for example the use of both sexes and inclusion of NGF in DRG neuron cultures which can alter chemosensitivity (Bevan & Winter, 1995). Thus far, the effects of NGF on factors related to IL-13 function in DRG neurons have not been investigated, but if, for example, IL-13Rα2 expression is upregulated, this would result in decreased neuronal stimulation by IL-13 due to its role as a decoy receptor (Khurana Hershey, 2003). A further indication that IL-13 is able to stimulate nociceptors arises from the finding that IL-13-sensitive neurons were frequently co-sensitive to capsaicin. These findings correlated with scRNA-seq analysis showing high levels of co-expression of TRPV1 and both IL-13 receptor subunits, IL-13Rα1 and IL-4Rα (Hockley et al., 2019). Whilst Ca^2+^ responses do not necessarily correlate with AP generation, these findings highlight the ability of IL-13 to stimulate sensory neurons and thus the potential for IL-13 modulation in conditions associated with visceral pain.

In TRPV1 knockout mice, IL-13-mediated activation of DRG neurons is reduced, thus implicating TRP channels in IL-13 signalling pathways (Oetjen et al., 2017). In support of previous findings, showing that the activation of p38 MAPK and JAK pathways is associated with TRPV1-mediated Ca^2+^ responses in cultured DRG neurons, we confirmed that simultaneous inhibition of JAK1, JAK2, JAK3 and Tyk2 by pyridone 6 reduced the population of IL-13-responsive neurons. More specifically, the effects of IL-13 on DRG neurons were regulated by JAK1/2, since blockade with ruxolitinib attenuated Ca^2+^ responses, recapitulating the effect of pyridone 6. Blockade of p38 MAPK also reduced neuronal stimulation by IL-13, in accordance with previous studies showing activation of p38 MAPK by IL-13 (Xu et al., 2003). Inhibition of the response to IL-13 by blockade of either p38 MAPK or JAK signalling may indicate an interaction between these pathways. Xu et al. (2003) made a similar observation wherein IL-13-induced STAT1/3 phosphorylation was attenuated by p38 MAPK inhibition.

Whilst the stimulation of DRG sensory neurons *in vitro* by IL-13 did not translate into direct changes in basal nerve firing *ex vivo*, IL-13-mediated sensitisation of colonic afferents in response to ramp distension was observed for the first time. Increases in afferent firing were noted at pressures >50 mmHg, implying that IL-13 sensitised high-threshold mechanonociceptors (i.e., mesenteric and serosal afferents) which correspond to a major class of nociceptor in the GI tract and comprise the majority of LSN afferent subtypes (Brierley et al., 2004). The mechanism through which IL-13 sensitises colonic afferent neurons is as yet unidentified, but somatically, in conditions with elevated levels of IL-13, sensory neuron TRP channel expression, including TRPV1, TRPA1 and TRPV4, is upregulated or the channels themselves are modulated to increase sensitivity (Feng et al., 2017; Moore et al., 2018; Wilson et al., 2013). Furthermore, in cardiomyocytes, IL-13 has been shown to modulate the voltage-gated Na^+^ channels Na_V_1.4 and Na_V_1.5 (Jude et al., 2019), but no work has thus far investigated if and how IL-13 modulates Na_V_ subunit expression in colonic sensory neurons (e.g., Na_V_1.8) (Hockley et al., 2019). Modulation of these ion channels may also occur in colonic afferent neurons to induce the hypersensitivity seen in response to noxious ramp distension, but further work is needed to understand precisely which channels are involved; for example, investigating whether IL-13 modulates expression or function of putative mechanosensors, such as Piezo2. Piezo2 is frequently co-expressed with IL-13Rα1 and IL-4Rα in lumbosacral DRG neurons innervating the colon (Hockley et al., 2019).

Multi-unit recordings of LSN activity showed that IL-13 did not sensitise capsaicin responses and therefore failed to modulate TRPV1 channels in colonic sensory afferents, suggesting other ion channels are responsible for afferent hypersensitivity. This is in contrast to TNFα which was found to sensitise the afferent response to capsaicin (Barker et al., 2022). IL-13 and TNFα interact with a similar population of TRPV1-expressing colon-innervating neurons and both seem to activate p38 MAPK (Hockley et al., 2019; Barker et al., 2022), so why does IL-13 not sensitise TRPV1? SB203580 (used in this study) inhibits both the α and β isoforms of p38 MAPK, which are highly co-expressed in colonic sensory neurons (Hockley et al., 2019). One may speculate that TNFα and IL-13 stimulate different isoforms of p38 MAPK, with TRPV1 being a substrate of only the p38 isoform activated by TNFα. TRPA1 has been linked to mechanosensitivity in the gut (Brierley et al., 2009; Cattaruzza et al., 2010) and, during inflammation, IL-13 increases TRPA1 expression (Oh et al., 2013). In scRNA-seq analysis, TRPA1 transcripts are present in a subset of TRPV1^+^ neurons, the same population that expresses IL-13Rα1 and IL-4Rα (Hockley et al., 2019). Further functional studies are required to investigate whether the IL-13-mediated colonic afferent hypersensitivity observed in this study is TRPA1-dependent.

Here, we have made the first report of p38 MAPK and JAK signalling in IL-13-induced mechanohypersensitivity in colonic afferent neurons. Previous studies have shown that IL-13 increases p38 MAPK phosphorylation (Xu et al., 2003) and JAK signalling has been assigned a role in somatic neuronal sensitivity and itch, in which IL-13 plays a key role (Oetjen et al., 2017). The distal colon is innervated by a range of sensory afferent populations, each with differing mechanical sensitivities (Brierley et al., 2004). Whilst these subtypes were not characterised in this study, an inhibitor of p38 MAPK signalling reduced afferent firing across all distending pressures (0-80 mmHg) and JAK inhibition affected afferent response at lower pressures (10-25 mmHg) and higher distending pressures (55-80 mmHg). These data suggest that p38 MAPK and JAK signalling pathways are involved in afferent responses to noxious mechanical stimuli, with serosal, mesenteric and muscular afferents likely those that are sensitised. However, ramp distension protocols only assessed afferent responses to stretch and, whilst this is most likely the predominant pain-evoking stimulus in GI disease, worsened by neuronal sensitisation, further characterisation of afferent subtypes sensitised by IL-13 may help to better understand the signalling pathways involved and provide novel targets for intervention. Data presented here also show that inhibition of p38 MAPK and JAK signalling had no effect on baseline nerve activity, demonstrating that normal neuronal function is not affected by blocking these pathways and thus may selectively target mechanisms of sensitisation to reduce pain in GI disease.

In summary, we have shown that IL-13 sensitises LSN responses to noxious ramp distension, an effect that can be reversed by inhibition of p38 MAPK and JAK signalling pathways. IL-13 receptor expression positively correlates with likely nociceptive populations of colonic afferent neurons (Hockley et al., 2019), making IL-13 and its receptors, alongside downstream p38 MAPK and JAK signalling, feasible targets in the treatment of visceral pain in GI disease.

## Additional Information

### Data availability

All data supporting the results presented in the manuscript are included in the manuscript

### Author Contributions

K.H.B. designed the research studies, conducted the experiments, acquired and analysed the data and wrote the manuscript. J.P.H. acquired and analysed the data and wrote the manuscript. L.A.P. analysed the data. I.P.C. designed the research studies. DCB, E.St.J.S. and FW designed the research studies and wrote the manuscript. All authors approved the final version of the manuscript submitted for publication and agree to be accountable for all aspects of the work in ensuring that questions related to the accuracy or integrity of any part of the work are appropriately investigated and resolved. All persons designated as authors qualify for authorship, and all those who qualify for authorship are listed.

